# SGFP-Grid : Split GFP Graphene Affinity Grids

**DOI:** 10.64898/2026.09.08.750207

**Authors:** Hanwen Feng, Jonathan Pickett, Qi Miao, Jiakai Hou, Weiyi Peng, Yimo Han, Zheng Zhou, Mingxing Teng, Zhao Wang

**Affiliations:** Verna and Marrs McLean Department of Biochemistry and Molecular Pharmacology, Baylor College of Medicine, Houston, TX, USA; Department of Pathology and Immunology, Baylor College of Medicine, Houston, TX, USA; Department of Biology and Biochemistry, University of Houston, Houston, TX, USA; Department of Materials Science and NanoEngineering, Rice University, Houston, TX, USA; Department of Molecular and Cellular Biology, Baylor College of Medicine, Houston, TX, USA; CryoEM Core (Advanced Technology Core), Baylor College of Medicine, Houston, TX, USA; Chemical, Physical, and Structural Biology Graduate Program, Baylor College of Medicine, Houston, TX, USA; Department of Molecular and Cellular Oncology, Division of Basic Science, The University of Texas MD Anderson Cancer Center, Houston, TX, USA

## Abstract

Affinity graphene grids provide a promising approach for selective protein capture in cryo-EM. Here, we introduce a split-GFP graphene grid platform(SGFP-Grid), in which graphene-conjugated GFP1–10 selectively captures GFP11-tagged proteins from low-concentration samples or cell lysates. This platform enables rapid assessment of target protein enrichment and particle distribution before vitrification via fluorescence imaging, while the grid design positions captured proteins away from the graphene surface and air–water interface. We also introduce a unique strategy to minimize nonspecific protein adsorption, thereby improving the selective enrichment of target proteins on this grid. Using GFP11-tagged apoferritin, we demonstrate fluorescence-guided protein capture and obtain a 2.58 Å cryo-EM reconstruction, establishing SGFP-G as an affinity-grid platform for high-resolution structural studies with reduced sample requirements.

## Main

Single-particle cryo-electron microscopy (cryoEM) has revolutionized the characterization of macromolecular assemblies that surpasses traditional methods. While advances in software and electron microscopes continue to be produced, sample preparation on grids for cryoEM has remained a bottleneck in obtaining data for high resolution structures^1,2^. Conventional single-particle cryoEM typically requires highly homogenous samples generated through protein overexpression and extensive affinity purification. These procedures can be time-consuming to optimize and may introduce sample degradation, loss of viable complexes, and batch variability. Functionalized affinity grids provide an alternative strategy by selectively capturing target proteins directly on the grid, particularly from samples in low concentration or endogenous proteins.

Affinity grids were introduced as early as the 1970s to address these challenges by functionalizing grid surfaces with ligands that selectively captured and immobilized selective plant viruses^3^. These types of grids were expanded further to include streptavidin crystal lipid monolayers for specific capture of biotinylated protein^4^. Subsequent developments introduced Ni-NTA lipid affinity grids for capturing His-tagged proteins^5^, nonfouling NTA-PEG coatings to reduce nonspecific adsorption^6^, and graphene oxide affinity grids incorporating the SpyTag/SpyCatcher system^7^. These affinity grid approaches reduced homogenous sample requirements and facilitated structural analysis of low-abundance or heterogeneous protein preparations.

Monolayer graphene grids are a complementary technology that can be used for functionalized cryoEM grids. In comparison to continuous carbon, graphene offers superior mechanical strength, atomic thickness, and improved signal to noise ratio^8,9^. Furthermore, monolayer graphene grids have been fabricated and shown to have improved cryoEM data for high resolution structural determination^10,11,12^. In addition, the surface chemistry of graphene can be modified to introduce functional groups for selective protein attachment and affinity capture^13,14,15^. Accordingly, several recent functionalization grid strategies have incorporated graphene/graphene oxide as the underlying support^7,16^. Together, these developments establish graphene as an attractive foundation for next-generation affinity cryoEM grids that combine low-background imaging with selective target enrichment.

Here, we report the design of a new split GFP graphene grid system capable of fluorescence detection to determine target particle enrichment on the graphene surface. The split GFP system is where the 11^th^ β strand is removed from the canonical GFP β-barrel to yield a pair of proteins (GFP 1-10/GFP11)^17^. The proteins do not emit green fluorescence independently and relies on the reassembly of the GFP β barrel shape (K_d_ = low μM to high nM) for fluorescence signal^18^. *In vitro* and *in vivo* models show the split GFP system is optimal for fluorescence microscopy detection, visualizing where the two proteins reassociate, and offer a system of high affinity reassembly ideal for proximity detection of two target proteins^19,20,21^.

We introduce the SGFP-Grid system where a graphene grid was conjugated with our novel GFP 1-10 on the grid surface and can capture low concentrations of GFP 11 tagged apoferritin (AP11). A high-resolution structure was determined using the low concentration of protein. This functionalized grid system offers a novel method of fluorescence detection not seen in previously developed technology, establishing a method to visualize protein particle enrichment before freezing in cryoEM sample preparation. We formed a complete pipeline of graphene grid production, functionalization, particle enrichment assessment, and application of a high-resolution single-particle cryoEM structure determination using SGFP-Grid.

## Results

### Design of SGFP-Grid

Monolayer graphene was added to a 1.2/1.3 holey carbon Au 300 mesh grid according to previously published methods^10^. The SGFP-Grid system involves the covalent linking of GFP 1-10 protein to the graphene surface (Figure 1). We first introduced defects to the monolayer graphene with ultraviolet light (UV) to add hydroxyl (-OH) chemical groups to the surface^22,23^. The hydroxyl groups were deprotonated in a reaction to prime them for reaction with carboxylic acid groups (-COOH)^24,25^. Carboxylic acid groups can react with free amine groups (-NH_2_) with the help of activating agents in an amine coupling reaction^26^. Hydroxyl groups on the graphene grid surface were functionalized with carboxylic acid. Detection using Toluidine Blue O (TBO)^27,28^ dye reveals our grid is functionalized with additional carboxylic acid groups (1.37 µM) compared to control graphene grid (0.213 µM) at 630nm absorbance (Figure 2A). The TBO detection of carboxylic acids establishes a way to quantitatively estimate the total number of functional group molecules on the surface relative to non-functionalized graphene.

**Figure 1.**
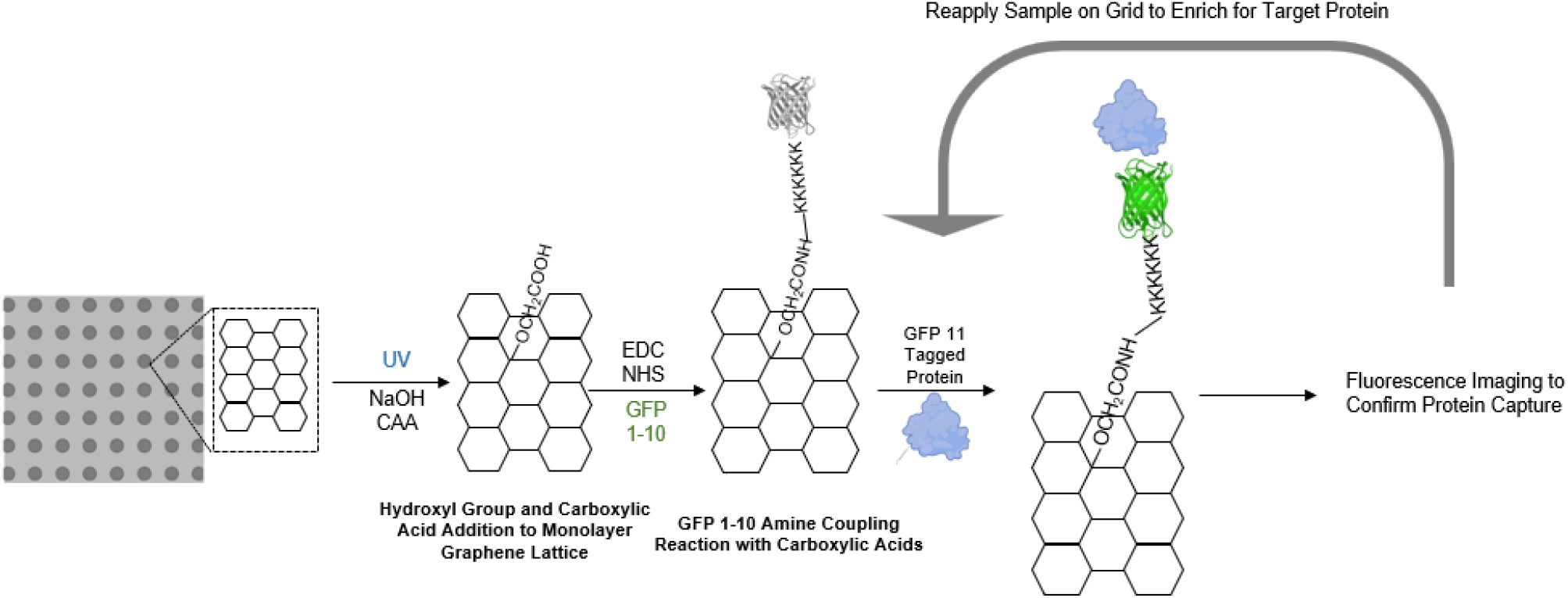
Functionalization of monolayer graphene grids with GFP 1-10 for affinity capture of GFP 11 proteins. Chemical conjugation of GFP 1-10 to monolayer graphene surface using UV/NaOH/CAA to add carboxylic acids followed by EDC/NHS to amine couple carboxylic acids to multi-lysine linker on GFP 1-10. GFP 11 tagged proteins, purified or from cell lysate, can be applied to the GFP 1-10 graphene grid with high affinity. Fluorescence of GFP 1-10 and GFP 11 reassembly *in vitro* will provide information on enrichment of target proteins on the grid surface. Multiapplication of protein sample can be performed until sufficient fluorescence is achieved. cryoEM grids can be prepared after obtaining structural information on target protein.

**Figure 2.**
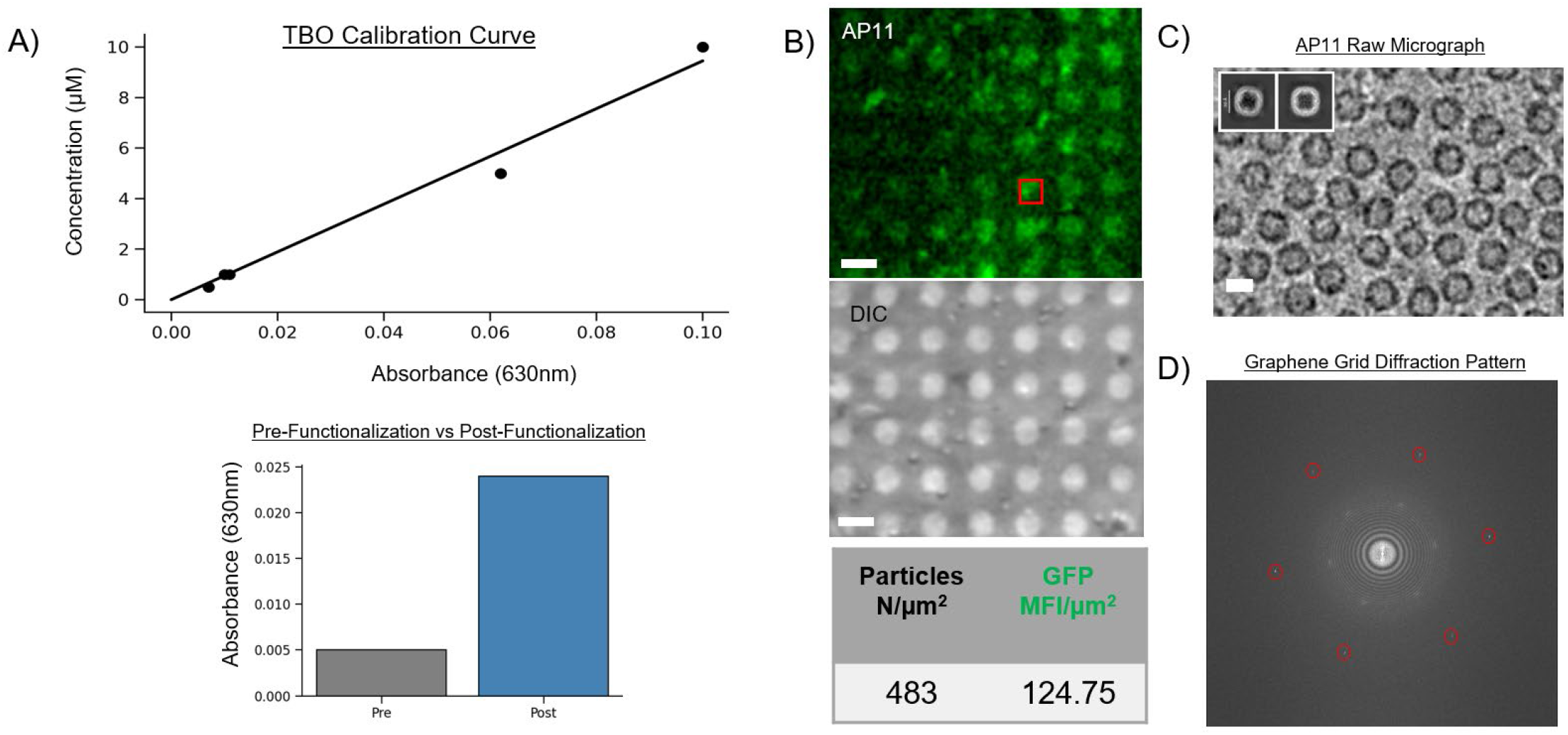
Carboxylation of monolayer graphene grids and the fluorescence of apoferritin GFP 11 (AP11) on GFP 1-10 graphene grids correlated with cryoEM particle distribution. A) Functionalization of graphene grid surface carboxylic acids is quantified with toluidine blue O (TBO). Concentration of carboxylic acids on the graphene grid surface increased after the functionalization steps (0.213 µM to 1.37 µM). Concentrations were calculated based on absorbance at 630 nm correlated to the concentration in µM of TBO/carboxylic acid. A) Fluorescence imaging of graphene covered grid square after 10x application of AP11 (0.005 mg/mL) to the grid. Green channel fluorescence and differential interference contrast (DIC) images are shown. Scale bar 2 µm. B) cryoEM micrograph showing the protein particle distribution embedded in vitreous ice. Diffraction pattern shows canonical six points representing the monolayer graphene lattice at around 2.2Å circled in red. Scale bar 10 nm. 2D classification of AP11 shows canonical 24mer cage structure capable of 3D reconstruction in cryoEM.

In the design of GFP 1-10 used in this affinity grid system, the final protein construct attaches to carboxylic acid groups on the graphene surface with a 6xlysine (6xK) N-terminal sequence (Figure 1). We fused a maltose binding protein (MBP) to the C-terminal to prevent GFP 1-10 aggregation into inclusion bodies when overexpressed in bacterial cells^29,30^. The purified mutant GFP 1-10 exhibited similar binding affinity and fluorescence in the presence of GFP 11 compared to a previously optimized (OPT) GFP 1-10^17^. We conjugated the mutated GFP 1-10 to the carboxylic acid groups on the graphene grid surface. Early testing reveals GFP 1-10 captured target protein in pairs (Supplementary Figure 3) potentially due to reported weak homodimer formation in GFP 1-10 *in vitro*^31,32,33^. Our final construct (Supplementary Figure 1) is the GFP 1-10 protein utilized in SGFP-Grid fluorescence imaging and capture of GFP 11-tagged target protein.

### Fluorescence Imaging Quantifies Particle Enrichment on Grid

To test successful graphene carboxylation and GFP 1-10 conjugation in the grid system, we designed and purified a GFP 11 tagged apoferritin (AP11) (Supplementary Figure 4) as a target protein to obtain a homogenous sample and control protein concentration. We tested particle enrichment capability of the SGFP-Grid system using extremely low concentration of AP11 (0.005 mg/mL) compared to cryoEM imaging conditions^1,2^. Our result demonstrates SGFP-Grid’s success in capturing AP11 at a concentration traditionally difficult to freeze grids. CryoEM screening image confirms AP11 has been captured and enriched on the grid (Figure 2C). The six-point diffraction spots seen in monolayer graphene was also visible in our high-magnification AP11 images which confirms graphene support film integrity^34,35^ (Figure 2D). Compared to other affinity grids, one advantage in the SGFP-Grid system is we can directly correlate fluorescence intensity to the enrichment of particle concentration on the grid. In our experimental condition, we identified number of AP11 particles (N=483) over a specific area (1 μm^2^) by cryoEM imaging (Figure 2B) and in parallel measure with fluorescence intensity (MFI) of ∼125 in fluorescence imaging. Based on fluorescent assay using purified mutants, we can correlate a certain number of molecules (N) per 1 μm^2^ of GFP 1-10 conjugated on the grid. Since our split GFP only generates fluorescence when it forms a complex with GFP 11, the fluorescence intensity can also indicate how much GFP 1-10 is occupied by GFP 11. SGFP-Grid can uniquely utilize fluorescence imaging to monitor whether the affinity grid is saturated by target protein.

The cryoEM data from affinity enriched AP11 yielded 2D classification with clear secondary features (Figure 2C). Following standard single particle processing, the 3D reconstruction of 24mer ∼500 kDa AP11 was processed to a 2.58Å resolution (FSC = 0.143) (Figure 3A). The final density map shows solid side chain fitting into the resolved density and low root mean square deviation compared to previously resolved apoferritin (PDB 8F49) (Figure 3B) (Extended Figure 6). The high-resolution map demonstrates there is no preferred particle orientation in the cryoEM data, which is supported by our design of GFP 1-10 containing a N-terminal flexible linker and AP11 containing 24 copies of GFP 11 extending from the 24mer assembly. The fully functionalized SGFP-Grid system establishes a relationship between GFP fluorescence and protein particles enrichment, as well as demonstrates capability of high-resolution 3D protein structure.

**Figure 3.**
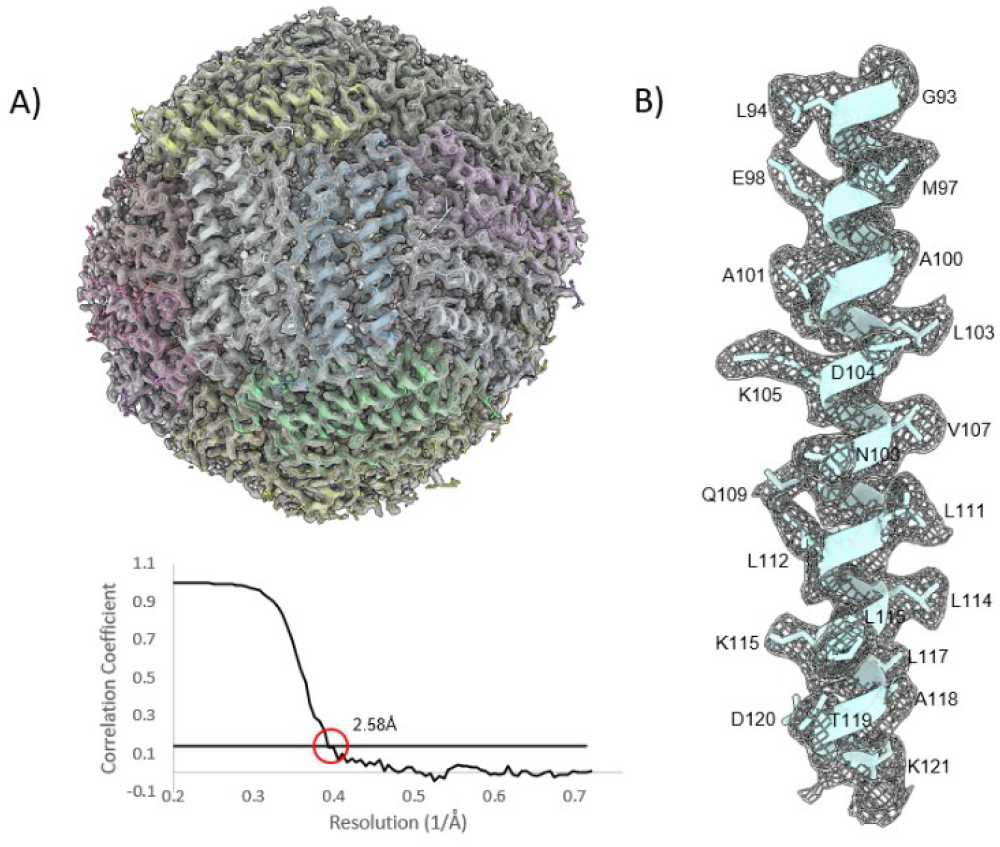
High-resolution cryoEM structure of GFP 11 tagged apoferritin (AP11). A) 2.58 Å AP11 structure shows clear secondary helical features in the density that directly matches the built model (FSC =0.143). B) Side chains of residues G93-K121 helix fit the cryoEM density with high resolvability.

### New Approach to Minimize Nonspecific Contaminated Protein on Affinity Grid

While using purified GFP 11 target protein confirmed the efficacy of SGFP-Grid to affinity capture proteins, affinity grids were established to bind target macromolecules in a heterogeneous sample^3^. Samples like cell lysates contain a complex mixture of proteins, nucleic acids, and lipids. Historically, graphene was avoided as material for grids due to its hydrophobicity and nonspecific contamination until a method was developed to change the surface properties of graphene to become hydrophilic and adsorb protein exceptionally well^36^. Therefore, samples containing contaminating proteins will nonspecifically associate with the graphene surface. In an affinity grid system, our goal was to enrich low abundance target protein and simultaneously minimize nonspecific binding from highly concentrated contaminating protein. Taking advantage of our SGFP-Grid system, we can easily detect enrichment of target protein and therefore set up a new adaptable strategy according to experimental conditions. Here, using recombinant mCherry protein, we mimicked an extremely low yield situation for our affinity grid. With mCherry being in high concentration and 1000x excess of target protein AP11 in concentration ratio, we performed fluorescent imaging first to detect changes in green and red fluorescence to optimize pipeline for high throughput.

All incubation and grid washing steps were performed at room temperature. Incubation of AP11 and mCherry mixture was performed on grids, with imaging of the grid after each iterative wash to detect fluorescence changes (Figure 4A). In our experiment, after 10x droplet sample incubation and over 30x dunking washes, mCherry signal decreases significantly (n=3, p<0.05) and represents the ability to wash away contaminating protein in high concentrations while GFP signal remains unchanged (Figure 4B). These results demonstrate that extensive washing can efficiently remove highly abundant nonspecific contaminants while retaining affinity-captured target proteins, establishing a fluorescence-guided strategy for minimizing nonspecific protein binding on SGFP-Grids.

**Figure 4.**
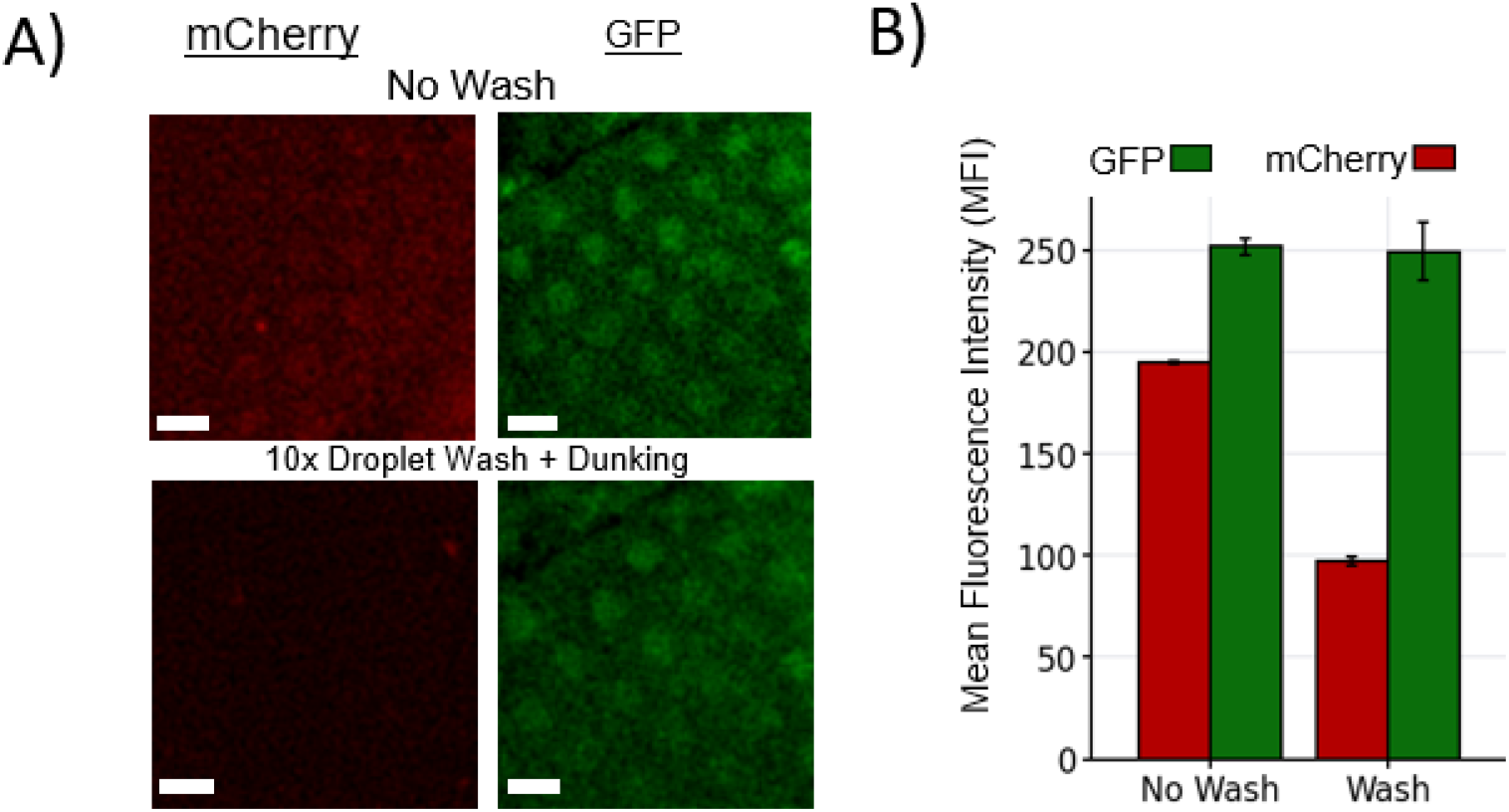
Quantification of contaminating protein removal via washing. A) mCherry (1000x excess) and Apoferritin GFP 11 (1x) was mixed and applied to the GFP 1-10 grid surface multiple times. Grid was washed in protein buffer. mCherry signal decreased visually while GFP signal remains similar intensity after 10x droplet washes + dunking in buffer. Scale bar 2 µm. B) Fluorescence was measured for each individual image (n=3). There was a significant decrease in fluorescence in mCherry fluorescence (p<0.05) while the average GFP fluorescence did not significantly decrease on the grid surface (p>0.05).

Overall, our SGFP-Grid system offers a novel approach to affinity capture GFP 11 tagged proteins, visualize protein particle enrichment at room temperature with GFP fluorescence, resolve a high-resolution AP11 structure captured by GFP 1-10, and enables a strategy for washing away of potential contaminating proteins at a ratio of 1000x times higher than target protein in a heterogeneous protein sample.

## Discussion

Functionalization experiment protocol on the graphene grid has been optimized to improve final particle distribution in our cryoEM results. Experimental use of other grids has not been tested as this is beyond the major purpose of our research design. Additionally, usage of other resources of graphene supply or different grid brands may slightly change these experimental conditions. Our final version of GFP 1-10 showed similar functional fluorescence as a previously optimized GFP 1-10 (OPT). Our mutated GFP 1-10 protein also demonstrated similar kinetics as the OPT GFP 1-10. In our biolayer interferometry (BLI) experiment, we identified a fast k_on_ and a slow k_off_ in our kinetic curves. These values represent a fast re-association of the GFP 1-10 and AP11 proteins, followed by slow release of the two bound proteins. Binding affinity also indicates a tight binding of GFP 1-10 and GFP 11 *in vitro* that surpasses other non-covalent and transient interactions. The identified behavior is beneficial to cryoEM grid preparation as it allows for successful fabrication of the GFP 1-10 grids and provides adequate time afterwards to successfully capture target proteins. Furthermore, we identified that mutating surface residues away from the chromophore center does not affect fluorescence and binding activity. Therefore, it remains possible to mutate GFP 1-10 further to identify mutations beneficial to *in vitro* binding between GFP 1-10 and GFP11.

In our study, we developed a GFP 1-10 affinity graphene grid using our mutated GFP 1-10 capable of capturing GFP 11 tagged proteins in low concentrations. Green fluorescence at room temperature before freezing demonstrates enrichment of protein particle on the grid surface. While other functionalized graphene grids offer affinity capture of target proteins, our system is the first to allow direct fluorescence imaging monitoring particle enrichment on grid before freezing. The fluorescence system allows for a faster approach to verifying protein particle enrichment on the grid surface in a non-cryogenic environment. This will be extremely useful for lowly abundant or non-steady expressed protein targets in the cell and enables monitoring of protein enrichment on the grid surface. Our 2.58Å structure of GFP 11 tagged apoferritin on our GFP 1-10 grids reveals the integrity of the protein on the grid surface is maintained after application, fluorescence visualization, washing, and freezing.

Monolayer graphene is shown to nonspecifically bind other protein and cellular components. We show that washing of the target enriched SGFP-Grid with protein buffer can remove our test nonspecific contaminated mCherry protein. mCherry was utilized in 1,000 X fold excess of apoferritin and represents contaminating protein in a sample mixture or cell lysate components. While nonspecific proteins associate with the grid surface, the binding affinity does not exceed the binding affinity between GFP 1-10 and GFP 11. Leveraging our unique grid system capable of utilizing fluorescent enrichment, our development of nonspecific contaminated protein washing procedures set us up to test with capturing GFP 11 tagged proteins directly from the cell lysate. It is noticeable that there are aggregates of mCherry that remain associated with the graphene surface even after extensive washing possibly because of mCherry denaturing and aggregating on the grid surface. Therefore, it is possible there are specific types of nonspecific binding to the graphene grid surface that will not be capable of being removed before freezing and require additional discovery in the future.

Our grids provide a streamlined approach to capturing a low abundance target protein with potential to skip protein purification processes that are time consuming and can lead to protein degradation before grid preparation. The SGFP-Grid can further expand the capabilities of current single-particle cryoEM to include endogenous proteins. Endogenous macromolecular assemblies are traditionally in low concentrations in the cell lysate and remain elusive to structural characterization. Genetic modification systems like CRISPR-Cas9 can assist in knocking in GFP 11 onto a target protein. Capture of the GFP 11 tagged endogenous proteins can then be performed using the GFP 1-10 graphene grids.

Overall, we introduced a new affinity graphene grid technology harnessing the split GFP system. Newly designed GFP 1-10 conjugated to the surface allow for capture of GFP 11 tagged target proteins, spontaneous reassembly of the GFP β-barrel structure, and chromophore maturation for fluorescence correlated with protein particle distribution on the graphene grid surface. We utilized our own mutated GFP 1-10 and could capture low concentrations of target protein and further wash away a highly concentrated contaminated protein in a heterogenous mixture. The capabilities of this system extend beyond low concentrations of purified proteins and into the eventual characterization of transient or endogenous proteins. This will ultimately translate to a deeper understanding of intermediate states of proteins and potential isoforms incapable of isolating using current biochemical purification methods.

## Methods

### Fabrication of Monolayer Graphene Grids

Monolayer grids were fabricated using 200 mesh 1.2/1.3 Quantifoil gold grids based on a previously published protocol^7^. Briefly, monolayer graphene on copper was purchased from Graphene Supermarket. PMMA Electron Beam Resist 495K (A4) was used to spin coat the graphene on a homemade spinner for 30s. The backside copper was etched using a plasma cleaner (80% argon/20% oxygen) and then removed using 1M ammonium persulfate. The graphene was scooped from the ammonium persulfate and left in distilled water for 20 minutes. Gold grids were used to scoop the monolayer graphene from the water and washed with acetone and isopropanol. The grids were then baked at 110°C for 15 minutes.

### Conversion of Graphene to Graphene Oxide on Grid Surface

Monolayer graphene grids were treated with UV light for 5 minutes. Then, the grids were incubated with sodium hydroxide and chloroacetic acid overnight with bubbling nitrogen gas. The grid was washed 3 times with Milli-Q water. The toluidine blue O (TBO) assay^27,28^ was performed on pre (UV only) and post (UV + carboxylic acid) functionalized grids to determine successful addition of carboxylic acid groups to the grid surface. 1mL of 5 mM TBO solution (pH 11) was incubated on the graphene grid surface for 10 minutes at room temperature. Then, after washing with base buffer (pH 11), bound TBO molecules to the surface was eluted using acidic base buffer (pH 2). The eluted volume was loaded onto a spectrometer to read the absorbance at 630 nm. A calibration curve was made using the original 5 mM TBO solution using a line of best fit (R=0.998). Concentration of the pre and post grids were calculated based on the calibration curve equation.

### Purification of GFP 1-10 and Apoferritin GFP 11 (AP11)

The GFP 1-10 mutated protein was fused to a maltose binding protein in a pMal-c5x vector and overexpressed in BL21 (DE3) *E. coli*. 5mL overnight cultures were used to inoculate 1L of LB media. Once OD_600_ reached 0.7, 0.5 mM of IPTG was added to induce recombinant protein production at 16°C overnight. *E. coli* were centrifuged and resuspended in purification buffer (40 mM Tris HCl, 200 mM NaCl, 1 mM DTT, pH 7.4). The bacteria were lysed by running through a microfluidizer press at 18,000 PSI 3 times. The lysate was centrifuged at 40,000 rpm for 1hr at 4°C to clarify and remove bacterial membrane components. The supernatant was incubated with amylose resin and rotated at 4°C for 1.5hr. The supernatant was allowed to flowthrough. The resin was washed with purification buffer and then incubated with D-(+)-maltose monohydrate to shake at 4°C for 1hr. The eluted protein was concentrated and loaded onto a Superose 6/150 size exclusion chromatography column using the base buffer (40 mM Tris HCl, 200 mM NaCl, 1 mM DTT, pH 7.4). The primary peak fraction was analyzed using SDS PAGE electrophoresis to confirm the successful purification of the fusion construct. The previously optimized GFP 1-10 (OPT) sequence was also cloned, expressed, and purified in a similar manner to the mutated GFP 1-10.

The human heavy chain of apoferritin was cloned into a pet30a vector with an N-terminal 6xHis tag and GFP 11. AP11 was transformed and overexpressed in BL21 (DE3) *E. coli* following a similar protocol to the GFP 1-10 construct. The AP11 pellet was suspended in GFP 1-10’s purification buffer with the addition of 20 mM imidazole, lysed with 18,000 psi 3 times, and clarified with 40,000 rpm for 1hr. The supernatant was removed and incubated with Ni-NTA resin for 2hrs rotating at 4°C. The supernatant was allowed to flowthrough, and the resin was washed with 150 mL wash buffer (40 mM Tris HCl, 200 mM NaCl, 1 mM DTT, 50 mM Imidazole, pH 7.4). The bound AP11 protein was then eluted with elution buffer (40 mM Tris HCl, 200 mM NaCl, 1 mM DTT, 300 mM Imidazole, pH 7.4). The AP11 protein was then ran through a Superose 6/150 size exclusion chromatography column using base buffer to ensure buffers match between GFP 1-10 and AP11. The homogenous peak was analyzed to confirm the successful purification of AP11.

### GFP 1-10 Functionality

The previously optimized GFP 1-10 (OPT) was also expressed in our MBP fusion system along with our mutated GFP 1-10 construct as a control. Both overexpressed proteins were purified using amylose resin followed by FPLC. Under different molar ratios of GFP 1-10 to AP11 (1:1/1:2/1:3), we performed a complementation assay. The assay was performed at room temperature and green fluorescence was detected using a plate reader. Both proteins showed significant improvement in fluorescence compared to empty well with only protein buffer (n=3, p<0.005) (Supplementary Figure 1B).

The biolayer interferometry assay was performed on an Octet. GFP 1-10, with a strep II tag, was first allowed to associate with the streptavidin sensor. After equilibration, GFP 1-10 was allowed to associate with various concentrations of AP11 (0.25 μM / 0.5 μM / 1 μM) for 300s. Then, the sensor with GFP 1-10/AP11 was moved to a well with only protein buffer for 300s after for disassociation

### Conjugation of GFP 1-10 to Graphene Grid Surface

GFP 1-10 was conjugated to the graphene grid surface using an amine coupling reaction. The carboxylic acids on the graphene grid surface are first activated using 2 mM n-(3-dimethylaminopropyl)-n’-ethylcarbodiimide hydrochloride and 5 mM n-hydroxysulfosuccinimide for 2hr at room temperature. Then, 0.35 µM of GFP 1-10 was incubated with the graphene grid overnight at 4°C. The grid was stored in buffer (40 mM TrisHCl, 200 mM NaCl, 1 mM DTT, pH 7.4) and stored at 4°C until usage.

### Fluorescence Imaging – Apoferritin GFP 11

The GFP 1-10 conjugated graphene grid was washed with protein buffer. Then, 5 µL of 0.005 mg/mL apoferritin GFP 11 was applied to the grid and incubated for 30s and hand blotted after. The application and blotting process was repeated 10 times total. Then, the grid was vertically dunked into protein buffer repeatedly for >30 times to wash away any nonspecifically bound proteins. The grid containing the apoferritin GFP 11 protein was imaged on a DeltaVision deconvolution microscope. Grid fluorescence were processed in ImageJ and DeltaVision software. Quantification of GFP fluorescence was done by subtracting the background grid signal from the targeted grid holes in triplicates in ImageJ.

### CryoEM Data Collection and Processing of Apoferritin GFP 11

516 raw movies were collected on a Titan Krios microscope equipped with a K3 detector with pixel size 0.68Å/pixel. The raw movies were motion corrected using MotionCor2^37^ in RELION^38^. The motion corrected micrographs were imported into cryoSPARC^39^ and CTF was estimated using CTFFIND4^40^. 120,000 total particles were template picked and final reconstruction of the particles using octahedral symmetry was processed to 2.58Å. The model of AP11 was built in ModelAngelo^41^ based on the final volume density. The final volume density was real space refined in PHENIX^42^ with the built model.

### Fluorescence Imaging – mCherry and Apoferritin GFP 11

mCherry was purchased from Origene (TP790040). mCherry (1000x) was mixed with AP11 (1x) and applied 10 times to the surface of the grid. The grid was imaged and then washed with 10 droplets of protein purification buffer before imaging again. Dunking washing involved using tweezers to keep the grid vertical and repeatedly dipping into protein storage buffer. The final fluorescence images were captured after this washing was performed.

## Acknowledgements

We thank T. Wensel for helpful discussion and feedback on the project design and fluorescence imaging. We thank the staff at UTHealth cryoEM core for the Titan Krios data collection for AP11 structure. We thank S. Aboulenaj for help in graphene grid fabrication.

## Contributions

H.F. and Z.W. conceived the project and designed the experiments. H.F. constructed the graphene grids, functionalized the grids, purified proteins, ran binding and fluorescence assays, and processed all data. Q.M. and M.T. assisted with initial carboxylation of graphene grids. J.P. and Z.Z. performed DeltaVision imaging for fluorescence images. Y.H. provided guidance on graphene grid fabrication. H.F. and Z.W. wrote the manuscript with input from all other authors.

## Ethics Declaration

H.F.,Z.W., and M.T. are contributors on BCM provisional patent application BLG 26-061.

